# The effect of proximity on the function and energy transfer capability of fluorescent protein pairs

**DOI:** 10.1101/838888

**Authors:** Jacob R. Pope, Rachel L. Johnson, W. David Jamieson, Harley L Worthy, Senthilkumar D. Kailasam, Husam Sabah Auhim, Daniel W. Watkins, Pierre Rizkallah, Oliver Castell, D. Dafydd Jones

## Abstract

Fluorescent proteins (FPs) are commonly used in pairs to monitor dynamic biomolecular events through changes in their proximity via distance dependent processes such as Förster resonance energy transfer (FRET). Many FPs have a tendency to oligomerise, which is likely to be promoted through attachment to associating proteins through increases in local FP concentration. We show here that on association of FP pairs, the inherent function of the FPs can alter. Artificial dimers were constructed using a bioorthogonal Click chemistry approach that combined a commonly used green fluorescent protein (superfolder GFP) with itself, a yellow FP (Venus) or a red FP (mCherry). In each case dimerisation changes the inherent fluorescent properties, including FRET capability. The GFP homodimer demonstrated synergistic behaviour with the dimer being brighter than the sum of the two monomers. The structure of the GFP homodimer revealed that a water-rich interface is formed between the two monomers, with the chromophores being in close proximity with favourable transition dipole alignments. Dimerisation of GFP with Venus results in a complex displaying ∼86% FRET efficiency, which is significantly below the near 100% efficiency predicted. When GFP is complexed with mCherry, FRET and mCherry fluorescence itself is essentially lost. Thus, the simple assumptions used when monitoring interactions between proteins via FP FRET may not always hold true, especially under conditions whereby the protein-protein interactions promote FP interaction.

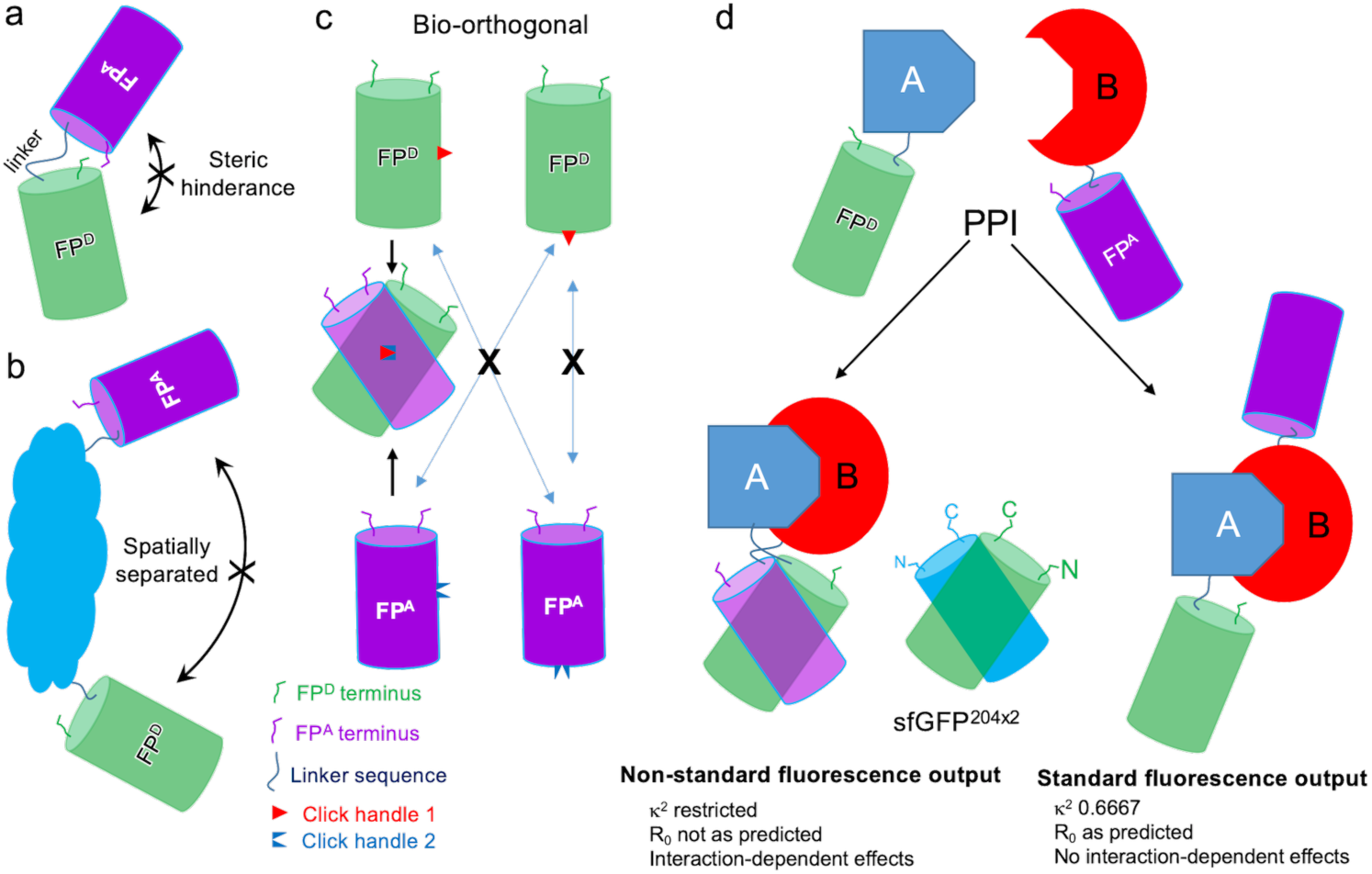

## Introduction

Fluorescent proteins (FPs) have revolutionised biology through their use as genetically encoded imaging tags and biosensors ^1-6^. The subsequent engineering of a small subset of natural FPs^1,2^, especially green fluorescent protein (GFP) from *Aequorea victoria* ^7^ and DsRed from coral ^5^ have expanded their use by changing their spectral (e.g. λ_max_, λ_EM_, quantum yield, brightness) and structural (e.g. quaternary structure, stability, folding kinetics, chromophore maturation kinetics) properties. One of the most important uses of FPs is to monitor dynamic biological events such as protein-protein interactions using processes such as Förster resonance energy transfer (FRET) ^8,9^. FRET is largely a passive process that relies on two FPs with mutually compatible spectral properties (acceptor FP absorbance overlapping with donor FP emission wavelength) being in close proximity; changes in distance between the two FPs changes efficiency of FRET between the donor and acceptor.

Despite FRET being a mainstay of biomolecular interaction analysis, there are a several assumptions required such as freely rotating FPs that do not interact or align in any significant manner. As well as absolute distance between the FPs, the angular vector between the chromophore dipoles is critical; this is the *κ*^*2*^ value equation 1.

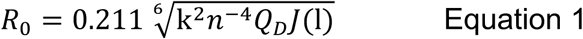

where *R*_*0*_ is the Förster radius, *κ*^*2*^ is the dipole orientation factor, *n* is the solvent refractive index, *Q*_*D*_ is the quantum yield of the donor and *J*(λ) is the overlap integral between the donor emission and acceptor molar absorbance. R_0_ is used as a constant to relate energy transfer efficiency to distance between individual components via equation 2.

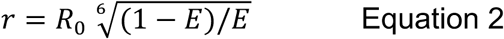

where r is the distance between two FRET chromophores and *E* is the observed FRET efficiency. Critically κ^2^ is arbitrarily set to 0.667 to reflect two randomly orientated chromophores as the dipole orientation is largely unknown which in turn impacts on the calculated *R*_0_. In reality the two chromophores are unlikely to be truly freely rotating with respect to each other when fused to a protein of interest ^9^. Therefore, it is difficult to accurately equate FRET efficiency to distance.

The question which thus arises is how the proximity of two FPs influence fluorescence, including FRET. Many fluorescent proteins, especially those that emit in the red region, naturally exist as oligomers ^10^ or have a tendency to oligomerise^11^. A great deal of protein engineering effort to generate functional monomeric forms but many commonly used FPs have been shown to have a capacity to dimerise ^11,12^. Dimerisation can be compounded by local high concentrations brought about by interactions between the fusion partner proteins that is to be monitored. Thus, when investigating FRET between FPs there may not just be simple spatial proximity at work but molecular interactions leading to more defined distance and dipole alignment, which may in turn influence inherent fluorescence. It has previously been thought that by using FPs from different organism classes with low sequence identities (e.g. GFP with RFPs) should prevent dimerisation.

We ^13^ and others ^14-17^ have previously shown that FP association can be promoted through either connecting FPs with linker sequences/protein domains, or by forming oligomers from individual monomers. In relation to the current work, we have shown that FP dimers can be constructed via genetically encoded strain-promoted azide-alkyne cycloaddition (SPAAC) ^13^, with dimerisation resulting in changes to the spectral properties. Here, we describe the construction and analysis of various Click linked FP dimers (Figure 1a). The structure of an artificial dimer of super-folder GFP (sfGFP) provides a rationale for enhanced fluorescence and role of water dynamics in this process. Using this new structural information, we determined κ^2^ values and measured *J*(λ) to calculate more realistic *R*_0_ values for experimentally analysed Click linked sfGFP-Venus dimers. We find that theoretical FRET efficiency does not match the observed FRET efficiency suggesting that proximity and dipole arrangement may not be the only factors that influence energy transfer. Furthermore, we linked sfGFP and mCherry together and found little FRET between the two proteins, with mCherry fluorescence being largely lost on dimerisation.

**Figure 1.**
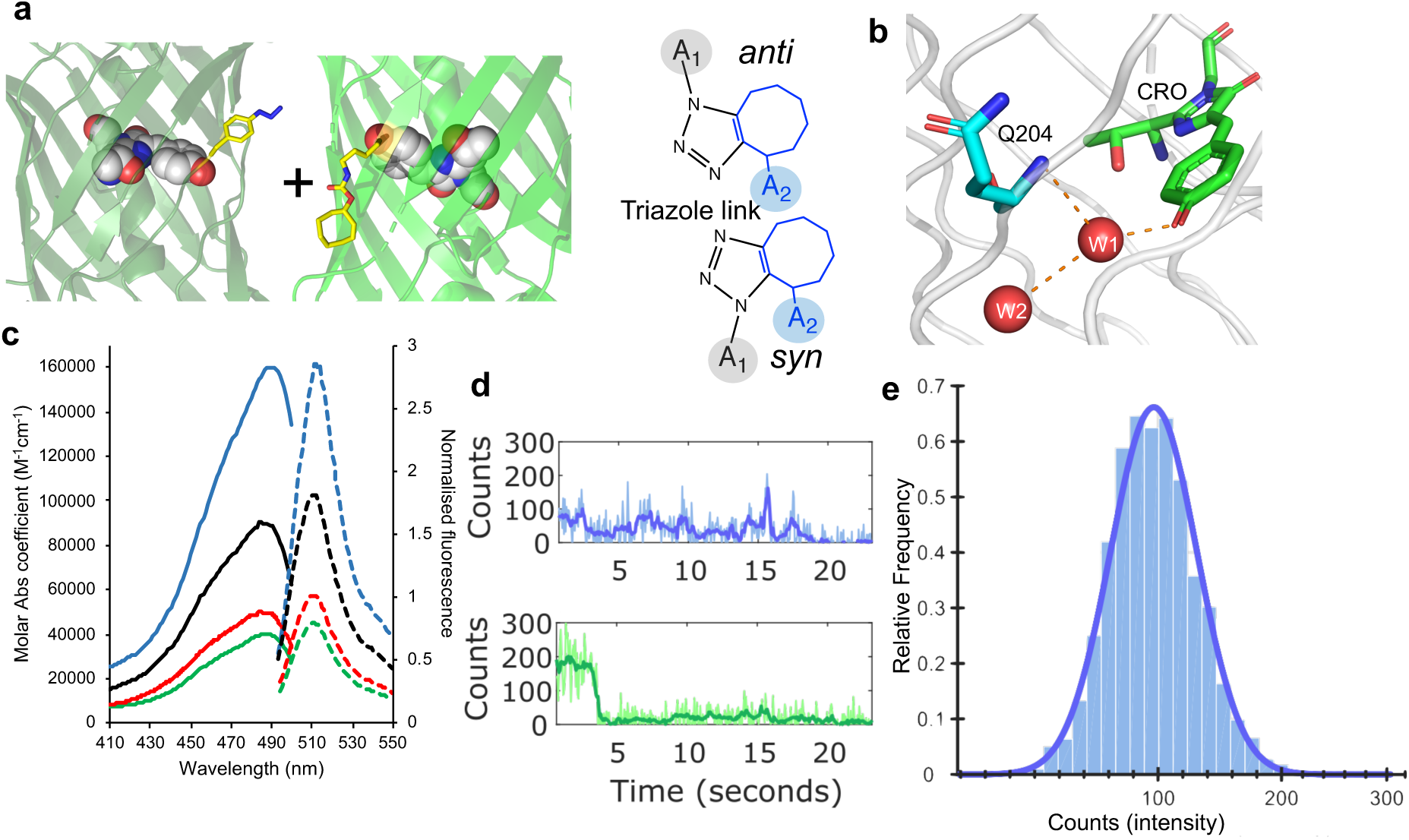
Click-based protein dimerisation via residue 204. (a) Covalent crosslink via genetically encoded *p*-azido-L-phenylalanine (azF) in one monomer and strained-cyclooctyne pyrrolysine (SCO-K) placed in the second monomer. Shown are the two different final regio-isomers available. (b) Relative positioning of residue Q204 with respect to chromophore (CRO) and local water molecules (red spheres W1 and W2). (c) Steady state bulk absorbance (full line) and fluorescence emission (dashed line) of sfGFP^204×2^ (blue), sfGFP^204azF^ (green), sfGFP^204SCO^ (red) and the addition of the two monomer spectra (black). The data has been reported previously ^13^ and shown here for context. (d) Representative single molecule traces for sfGFP^204×2^ (blue) and sfGFP^WT^ (green) measured by TIRF microscopy. Further example of single molecule traces can be found in Supporting Figure S1 for sfGFP^204×2^ and work by Worthy et al ^13^. (e) A single molecule fluorescence intensity histogram for sfGFP^204×2^ consisting of 179 trajectories (2602 spots). The histogram data fits to a single log normal distribution centred around 100 counts.

## Results and discussion

### The effect of sfGFP proximity on function

We have previously reported the construction of artificial FP dimers by Click chemistry through the covalent coupling of genetically encoded of ring-strained cyclooctyne derivative of the pyrrolysine (SCO-K) and *p-*azido-L-phenylalanine (azF) ^13^ (Figure 1a). It should be noted that we do not attempt to change residues at the FP dimer interface nor link them in a tandem arrangement using a spacer sequence as has been done in other approaches ^14-18^ but model potential naturally occurring interface sites, which are in turn stabilised through a SPAAC link. Regions that do not naturally associate do not promote covalent crosslinking via SPAAC ^13^. Thus, our approach stabilises naturally feasible protein interactions.

Residue Q204 in sfGFP lies close to the chromophore (CRO; Figure 1b), with the backbone amine group making an indirect H-bond with CRO via a conserved structured water molecule, W1. *In silico* molecular docking revealed that Q204 consistently resided at possible dimer interfaces and is close to a region known to be involved in FP dimerisation ^12^. The SCO-K (sfGFP^204SCO^) and azF (sfGFP^204azF^) containing monomers were subsequently proved to dimerise, generating the dimer termed sfGFP^204×2 13^. The sfGFP 204-linked dimer displayed enhanced fluorescence compared to the monomers. Dimeric sfGFP^204×2^ displayed positive functional synergy in which the brightness of the complex was more than the sum of the individual monomers (Figure 1c) ^13^. Indeed, sfGFP^204×2^ is brighter on a per CRO basis (57000 M^-1^cm^-1^) than the original sfGFP (37000 M^-1^cm^-1^) ^19,20^ and EGFP (33000 M^-1^cm^-1^) ^21^, two benchmark fluorescent proteins.

The increase in molar absorbance coefficient suggests that the fluorescence lifetime is shorter for sfGFP^204×2^ (0.92ns) compared to the original monomeric sfGFP (3.2 ns, calculated using the website huygens.science.uva.nl/Strickler_Berg/). Real time single molecule fluorescence of sfGFP^204×2^ was undertaken to explore the mechanism of enhanced capacity of the dimer to absorb and emit light. The dimer has an increased ON time compared to the sfGFP^WT^ (average 0.87s GFP^204×2^ compared to 0.65s for GFP^WT^). Analysis of individual traces shows this clearly as sfGFP^204×2,^ takes longer to photobleach compared to sfGFP^WT^ (Figure 1d with additional traces in Supporting Figure 1; also see Worthy et al for WT sfGFP single molecule analysis^13^). The increased ON times and photobleaching lifetime coupled with the shorter fluorescence lifetime is likely account for the increased fluorescence observed in steady state ensemble measurements. It is notable that the single molecule fluorescence time course traces are more complex, and dynamic compared to sfGFP^WT^ with a range of fluorescent states observed, which could indicate cooperative interaction between the individual monomer units.

Ensemble histograms reveal a single dominant intensity peak is observed at a value equivalent to monomeric sfGFP^WT^ (Figure 1e). This differs from a previously described artificial dimer linked via residue 148 (termed sfGFP^148×2^) ^13^, which exhibits two distinct population states. If the two molecules in the dimer are acting largely independently of each other, a bimodal distribution would be expected. Thus, only 1 CRO in the dimer is fluorescent at any given time.

### Structural basis for proximity-based effects

The structure of sfGFP^204×2^ (structural statics in Supporting Table S1 and Supporting Figures S2a-b) reveals that each monomer unit is similar to the original starting sfGFP. The sfGFP^204×2^ dimer forms a quasi-symmetrical off-set “side-by-side” monomer arrangement (Figure 2a), which is promoted by formation of a *syn* 1,5 triazole link that generates a reverse turn structure (Figure 2b). The two CROs points towards each other in an antiparallel arrangement 22 Å apart with a 5Å offset (Figure 2c). It is closest to the 3rd ranked model predicted previously ^13^ (Supporting Figure S2b-c). Each monomer is offset by ∼70° with the C-termini close in space (Figure 2b). As the N- and C-termini are close to each other at the same end of the β-barrel, the proximity and orientation of the two termini in the dimer may well promote such an interaction in a fusion protein construct.

**Figure 2.**
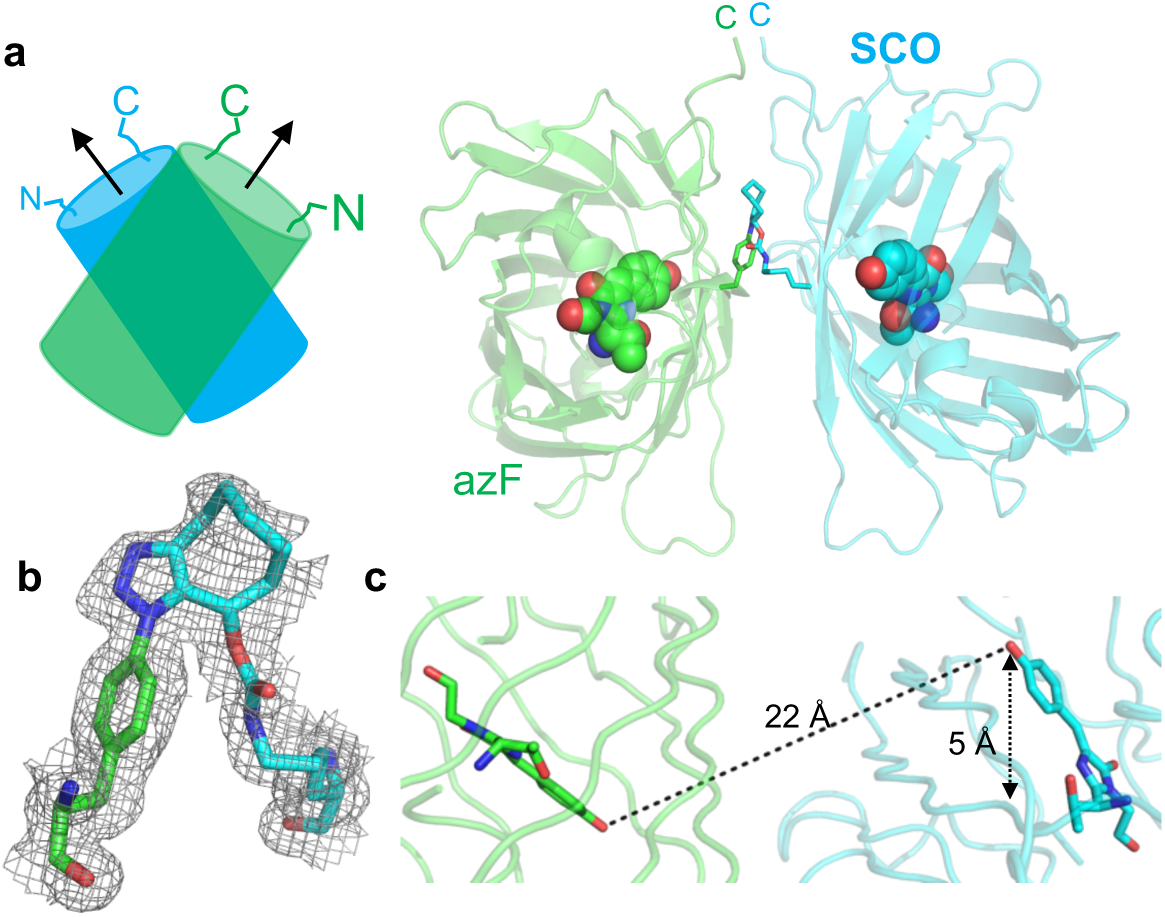
Structure of sfGFP^204×2^. (a) Arrangement of the azF (green) and SCO (blue) containing monomers. (b) The *syn* conformation of the triazole linkage with the electron density map (2Fo-Fc, 1.0 sigma) shown. (c) Distances and offset of the two CROs (shown as sticks)

The two monomer units associate to form an extensive and intimate interface. While the interface area is relatively small (∼900 Å^2^), the main elements that comprise a protein-protein interface, namely hydrophobic interactions and H-bonding are observed (Figure 3). The H-bond network at the interface is not symmetrical but the hydrophobic interactions show a significant degree of symmetry (Figure 3a). The hydrophobic core interface is comprised of Phe223, Val206, Leu221 from both chains interlocking (Figure 3b). These residues are surface exposed in sfGFP and form a naturally occurring hydrophobic patch ^12^ that can facilitate and stabilise the dimer on Click crosslinking (Figure 3e), or for that matter potentially other FPs. Indeed, mutation of Val206 to a charged residue is known to reduce dimerisation tendency of *A. victoria* derived GFPs ^12^.

**Figure 3.**
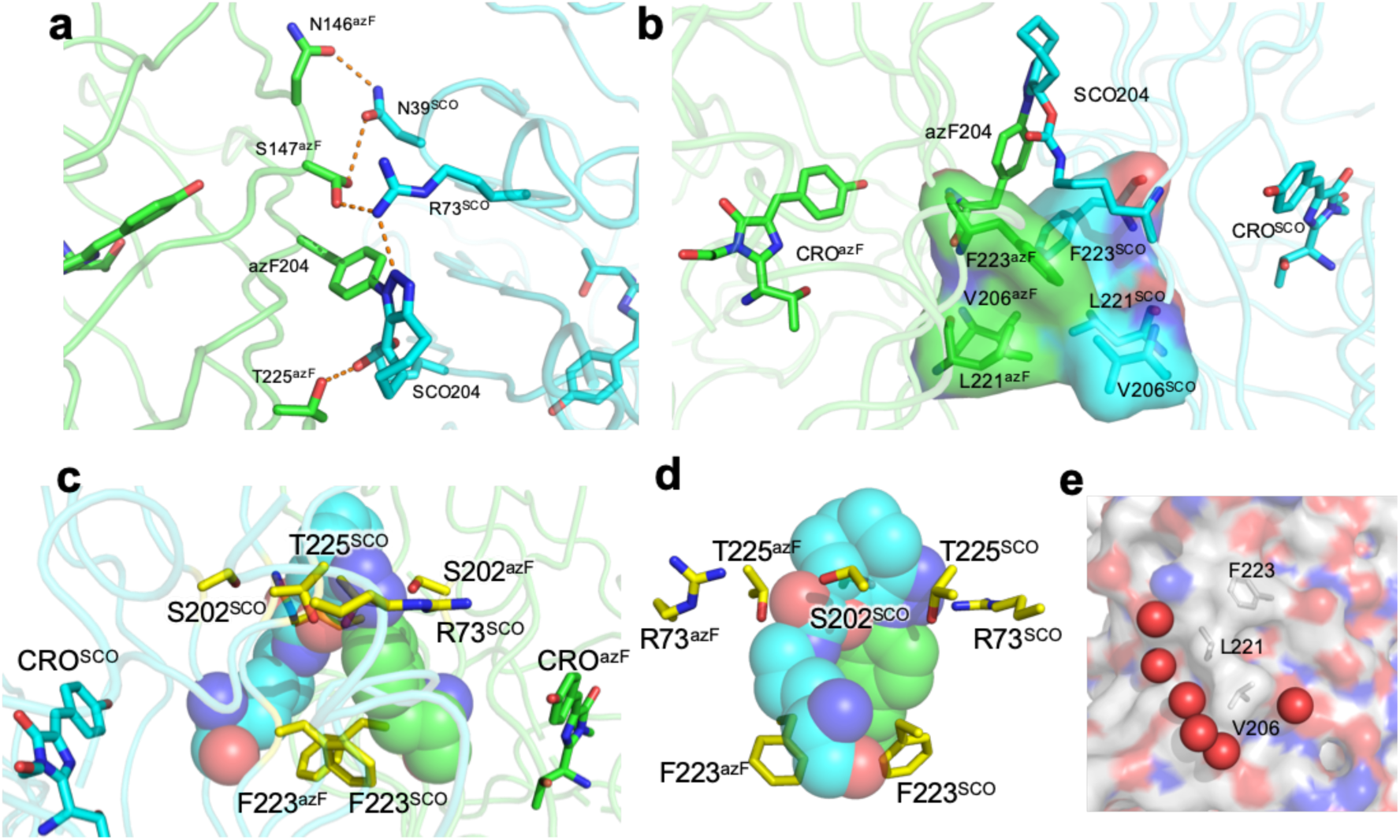
Subunit interface in sfGFP^204×2^ comprised of sfGFP^204azF^ (green) and sfGFP^204SCO^ (cyan). (a) H-bond network at interface. (b) Hydrophobic interactions. (c and d) Interactions around the triazole link shown in two different orientations. (e) Water molecules (red spheres) associated with the interface region.

The new triazole crosslink is integrated within the structure being semi-buried at the dimer interface and lying above the plane of the main hydrophobic interface patch (Figure 3c-d). The azF component is fully buried while one face of the SCO moiety is partially accessible to the solvent. Phe223 from both monomers forms the base of the triazole reverse turn (Figure 3c-d) while Arg73, Ser202 and Thr225 residues make putative polar interactions with oxygen and nitrogen atoms in the SCO-azide link. A more extended network linking the two chromophores is proposed in Supporting Figure S3. Thus, the new crosslink is not just a simple chemical bolt link between the two monomers but forms an integral structural component.

At the interface are two cavities filled with ordered water molecules (Figure 4a). The water molecules are arranged around an area where the chromophore protrudes towards the surface. A partially buried water molecule (W1; Figure 1c and Supporting Figure S4) is commonly observed associated with the chromophore via a H-bond with the hydroxyl group and the backbone of residue 204; this water is associated with 1 to 2 additional surface water molecules (grey spheres, Figure 4b-c) as observed for monomeric sfGFP^WT^ (Supporting Figure S4). In the sfGFP^204×2^ dimer, these waters lie within the cavity together with several additional tightly packed water molecules. The roles of the additional waters associated with W1 in terms of their impact on the structure-function relationship is not fully known but it has been postulated that they contribute to charge transfer and modulating the protonated state of the CRO ^22-24^. In solution, it is likely that the additional water molecules associated with W1 are in free exchange with the solvent when sfGFP is monomeric; exchange with bulk solvent is likely to be minimal in the dimeric sfGFP^204×2^ so persist in a defined arrangement for longer. By changing the dynamics of normally surface associated water molecule could potentially contribute towards the enhanced brightness observed on dimerisation through the formation of more persistent networks.

**Figure 4.**
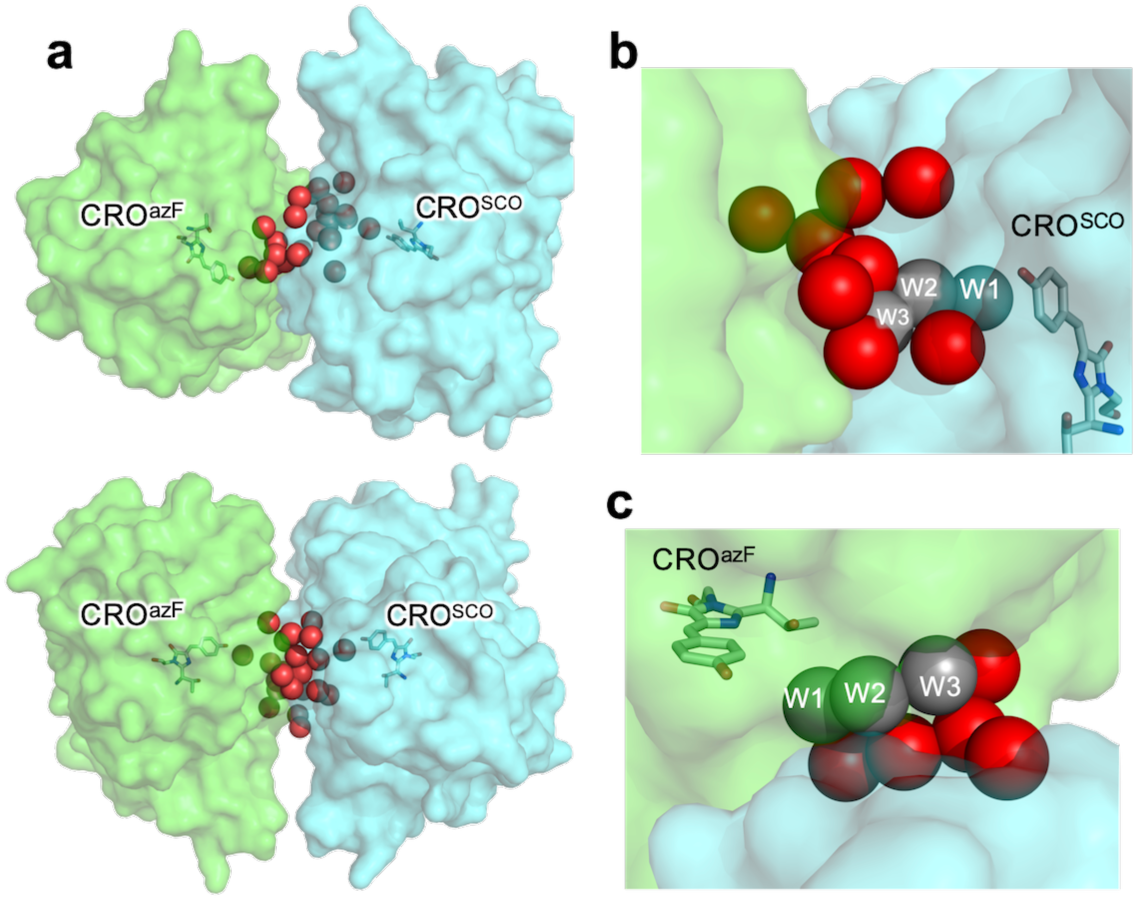
Water-rich cavities at dimer interface. (a) water (red spheres) filled cavities shown from two different angles. Waters associated with the (b) sfGFP^204SCO^ CRO and (c) sfGFP^204azF^ CRO. The grey spheres are equivalent to W1, W2 and W3 shown in Figure 1 and Supporting Figure S4. Waters molecules W2 and W3 are observed in the sfGFP^WT^ structure but are largely surface exposed (Figure S4).

### Heterodimers and functional communication by energy transfer

The use of different FPs with compatible spectral properties to promote FRET is essential for biomolecular analysis. The sfGFP^204SCO^ variant can be linked to Venus (containing azF) via residue 204 to generate heterodimers ^13^. The resulting dimer, termed GFVen^204^, demonstrated FRET from the sfGFP component to Venus, as would be expected (Figure 5a). There is currently very little known about the relative orientation of FRET-based FP pairs with only one structure available in a biosensor configuration ^14^, which is in a single polypeptide format rather that a classical two-protein system. Given the high degree of sequence and structure similarity between sfGFP and Venus, we used the GFP^204×2^ structure to build models of the GFVen^204^ dimer so as to calculate more specific *R*_0_ factors based on the relative orientations of the two chromophores (Supporting Figure 5). Using our model of GFVen^204×2^ together with the known transitions dipole arrangements for both GFP and Venus ^25,26^ (Figure 5b), *κ*^*2*^ was calculated in the model to be 3.59. Using the Q_D_ and *J*(λ) values (Supporting Table S2) together with a refractive index of 1.4 to account for a combined protein-water environment (Hellenkamp et al ^9^ and Dr Tim Craggs personal communication via Twitter) we calculated *R*_0_ with the different κ^2^ values (Supporting Table S2). The calculated R_0_ differs were ∼76 Å, which is up to 19 Å longer when calculated using the arbitrary 0.667 κ^2^ value. Our calculated R_0_ values are consistent with those calculated using *J*(λ) and donor QY values available through FPbase (https://www.fpbase.org) ^27^ when adjusted for κ^2^ (see Supporting Table S2).

**Figure 5.**
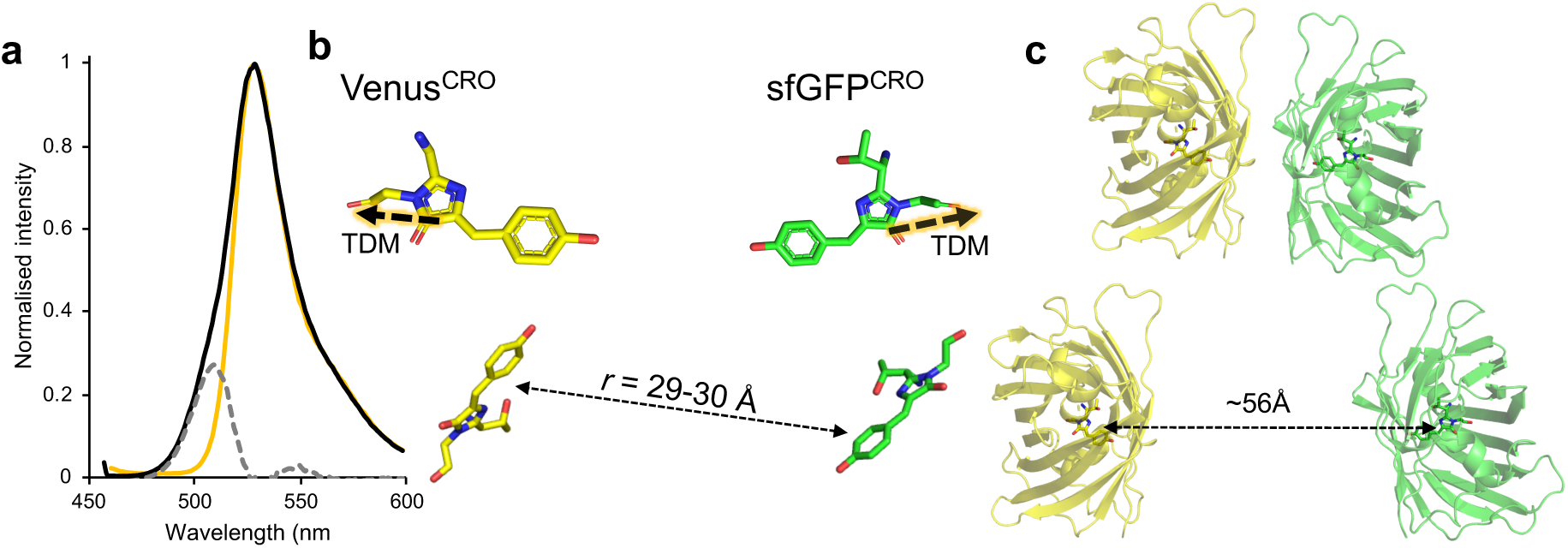
Energy transfer between sfGFP and Venus. (a) Fluorescence emission spectra of GFVen^204^ (black), Venus^204azF^ (gold) and the residual emission profile highlighting sfGFP contribution to GFVen^204^ spectrum (grey dashed). Excitation was at 450 nm. (b) Relative positionings of the Venus (yellow) and sfGFP (green) chromophores (CROs) in the model GFVen^204^ structure. The dashed lines highlighted in orange represent the transition dipole moment (TDM). (c) Relative distances between Venus (yellow) and sfGFP (green) based on the model of SPAAC linked dimer (top) and observed FRET efficiency (bottom). Structures are to scale.

The question arises is how does our calculated *R*_0_ relate through to observed FRET efficiency. Based on the use of equation 2 and the measured inter-chromophore distance of 29-30 Å (Figure 5b), the estimated FRET efficiency for our GFVen^204^ construct should be close to 100% (99.6-99.7%). However, deconvolution of the GFVen^204^ emission on excitation at 450 nm (a wavelength that will excite only sfGFP) reveals a significant sfGFP component (Figure 5a). Using equation 3, FRET efficiency can be calculated.

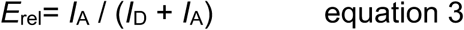

where *E*_rel_ is relative FRET efficiency, *I*_A_ is integrated fluorescence of the acceptor and *I*_D_ is the integrated fluorescence of the donor. FRET efficiency was determined to be 86.8%. Thus, there is a clear discrepancy between the observed and theoretical FRET efficiency, which has been observed before form structure-based analysis where inter-FP interactions were observed ^7,14^. What gives rise to this difference? A simple and obvious explanation is that some free monomeric sfGFP^204SCO^ is present. Analysis of polyacrylamide gels and mass spectrum suggests little or no monomeric protein is present (see Worthy et al ^13^ and Supporting Figure S6 for details). Are the considerable number of water molecules present at the domain interface observed for sfGFP^204×2^ (Figure 4) playing a role in quenching? Water can quench fluorescence ^7,28^, especially if collisional events are promoted through free dynamic exchange. However, the crystal structure suggests local water molecules are likely to be less dynamic in the dimer compared to monomeric forms. Is the arrangement of the monomers in GFVen^204×2^ similar to the assumed sfGFP^204×2^? While we cannot rule out some rotation of one FP with respect to another, the triazole link will restrict such rotation and the CROs will retain a similar vector configuration in terms of the transition dipole moments. With a R_0_ of 76.62 Å, the two CROs will need to be at least 50 Å apart (shown schematically in Figure 5c). Even using the arbitrary κ^2^ value of 0.667 generates a R_0_ of 56 Å, which will require the CROs to be ∼40 Å apart to generate the observed FRET efficiency. Given the relationship of residue 204 to the CRO (Figure 1), neither distances are feasible in a covalently linked dimer. It is clear that bringing two different FPs in close proximity so promoting inter-FP interactions can influence FRET efficiency, which results in an overestimation of the distance between the pair.

### Proximity effect of green and red fluorescent proteins

We next linked together sfGFP with a DsRed derived monomeric protein, mCherry ^30,31^. Green fluorescent proteins can be used as a FRET partner with mCherry^16,32-34^ with an estimated *J* coupling of 1.8×10^15^ M^-1^cm^-1^nm^4^ (FPbase FRET tool (www.fpbase.org/fret/) ^27^. The sfGFP^204SCO^ variants was reacted with mCherry containing azF at the structurally equivalent position, residue 198 (Figure 6a). Molecular docking suggested the two proteins can associate at the interface between residues 204^sfGFP^ and 198^mCherry^ (Figure 6a), with covalent coupling via SPAAC subsequently proved by SDS PAGE (Supporting Figure S7). Incorporation of azF at residue 198 in mCherry had little effect on the spectral properties of the momomer with a similar molar absorbance and brightness to the wt mCherry (69,000 M^-1^cm^-1^ with a quantum yield of 24% compared to 72,000 M^-1^cm^-1^ for wt mCherry with quantum yield of 22% at 587 nm; Figure 6c and Supporting Figure S8a).

**Figure 6.**
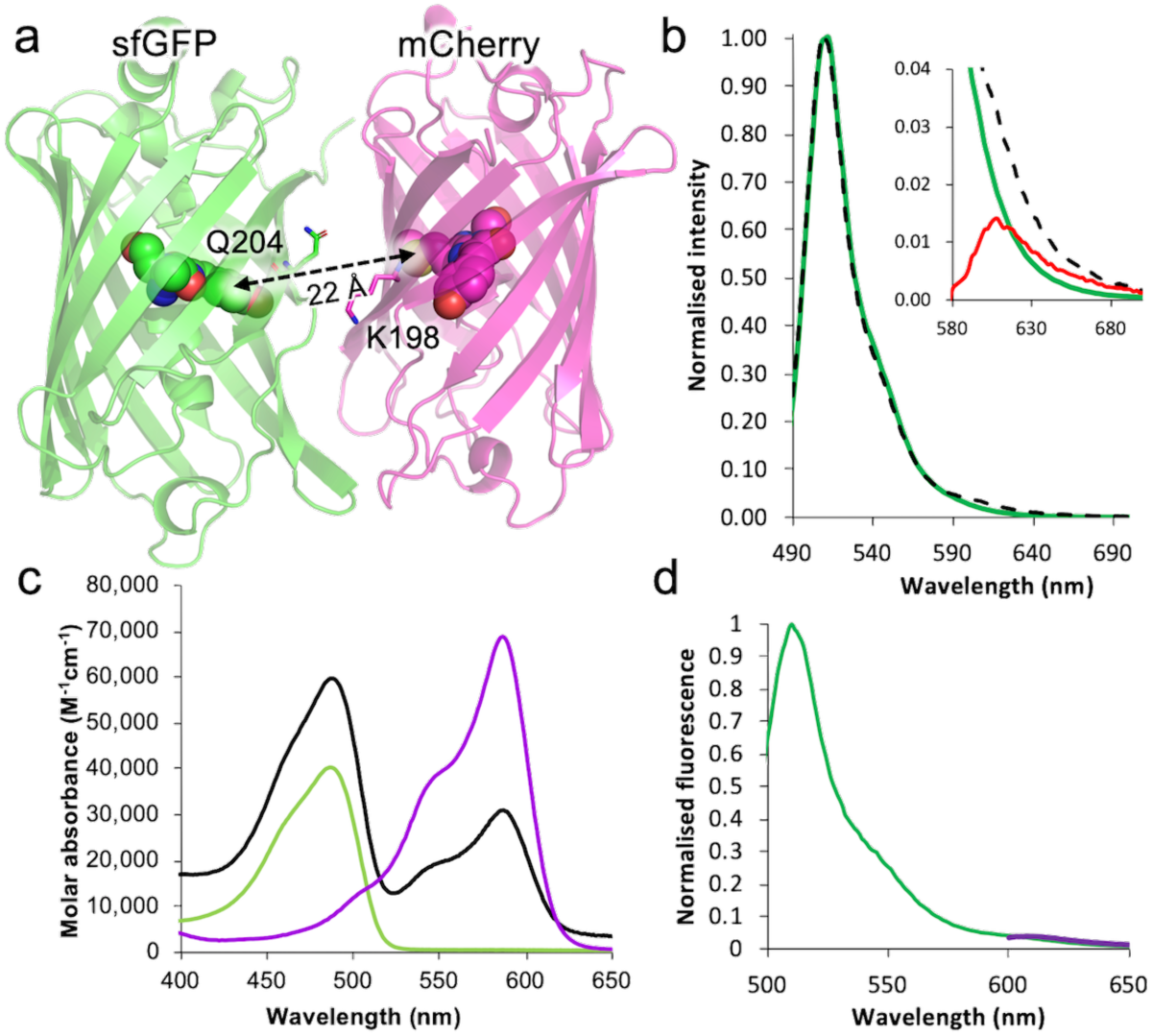
Effect on function of proximally located sfGFP and mCherry. (a) Modelled docking of mCherry (PDB: 2h5q) and sfGFP (PDB: 2b3p) using ClusPro^29^. Residues Q204 (sfGFP) and K198 (mCherry) that are replaced by SCO-K and azF, respectively, are highlighted. (b) Fluorescence emission (on excitation at 485 nm) of sfGFP^204SCO^ (green solid line) and GFCh^×2^ dimer (dashed black line). Emission intensities are normalised to sfGFP^204SCO^. Inset is the zoom in region of the emission spectrum centred around 610 nm, the emission maximum of mCherry. The red corresponds to the subtraction of GFCh^×2^ from sfGFP^204SCO^. (c) Molar absorbance of sfGFP^204SCO^ (green), mCherry^198azF^ (purple) and GFCh^×2^ (black line). (d) Emission profile of GFCh^×2^ on excitation at 485 nm (green) and 585 nm (purple).

The purified dimer, termed GFCh^×2^ did not appear to display any significant FRET on excitation at 490 nm (Figure 6b). Indeed, very little observable fluorescence can be attributed to mCherry in the dimer even on excitation at 585 nm (Figure 6d), which is confirmed visually through general UV excitation (Supporting Figure S7b). The mCherry associated peak at ∼585 nm is reduced in terms of molar absorbance compared to the mCherry^198azF^ monomer. As with other dimeric forms (*vide supra* and ^13^), the sfGFP molar absorbance increased above the simple addition of the two monomeric forms (∼16,000 M^-1^cm^-1^ taking into account the contribution from the mCherry chromophore) confirming the role of dimerisation via residue 204 in enhancing sfGFP function. Data suggests that the interaction between the sfGFP and mCherry is responsible for loss of fluorescence. Reaction with the SCO-K ncAA alone does not appear to affect fluorescence (Supporting Figure S8b). Attachment with the bulker azide containing Cy3 dye also does not result in loss of fluorescence, with FRET observed as expected (Supporting Figure S8c). Thus, placing FPs in close proximity to promote their interaction is the likely course of the loss in fluorescence, in this case from mCherry. As donor fluorescence is still observed, in a FRET experiment this could be interpreted as two proteins not interacting when the opposite may in fact be the case.

### Comparison with alternative sfGFP dimer sfGFP^148×2^

The structure of another Click-linked artificial dimer joined via residue 148 (termed sfGFP^148×2^) has recently been reported ^13^. Dimerisation effectively switched sfGFP^148×2^ on, with the dimer displaying improved function compared to both monomers and the original wild type sfGFP (sfGFP^WT^). We used the structure of the sfGFP^148×2^ dimer to calculate κ^2^ as a representative alternative CRO arrangement. This will in turn allow us to investigate how different configurations of one monomer to the other affect dipole alignments and hence FRET. Residue 204 lies close to 148 on the adjacent β-strand (Figure 7a) but they adopt very different sidechain and thus monomer arrangements in their crystal structures (Figure 7b-c). In contrast to sfGFP^204×2^, the triazole link in sfGFP^148×2^ forms the extended *anti* form that is re-enforced with both polar and hydrophobic interactions between the monomers generating a quasi-symmetrical “head-to-tail” arrangement the monomers. The result of such a configurational change between the two monomers units results in the relative positioning of the CROs being very different (compare Figure 7b and 7c).

**Figure 7.**
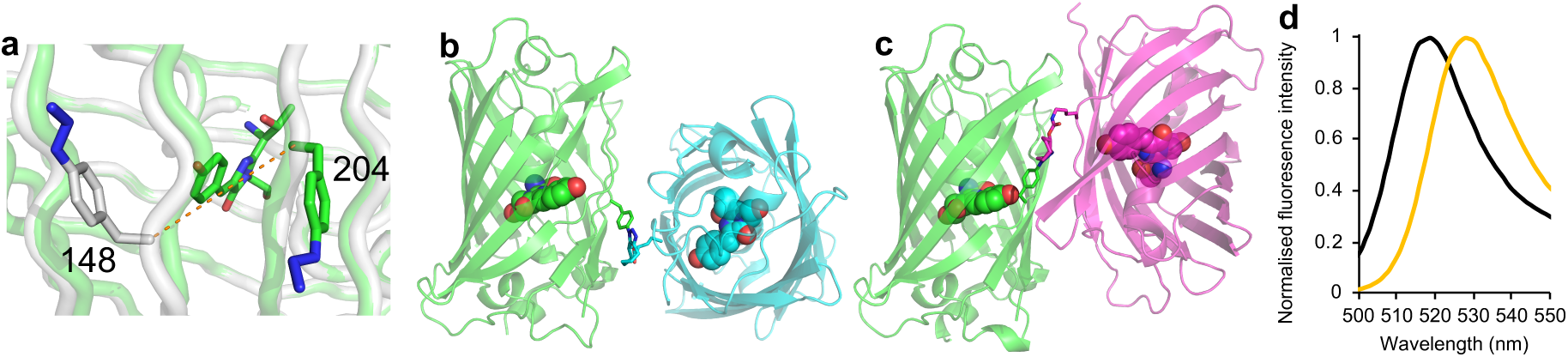
Comparison of sfGFP^204×2^ structure with sfGFP^148×2^. (a) Location of the azF moiety at residue 148 (grey) or 204 (green). Structure of (b) sfGFP^204×2^ and (c) sfGFP^148×2^. Each structure is orientated identically with respect to the azF containing ncAA (coloured green) to highlight the relative differences in monomer arrangements. (d) Normalised emission of on excitation at 490 nm for GFVen^148^ (black) and wt Venus (gold).

Using the same approach as for GFVen^204^, we calculated κ^2^ values for a model of GFVen^148^ based on the sfGFP^148×2^ structure. It should be noted that unlike linkage through residues 204 (or 198 in mCherry), covalent coupling via residue 148 was designed to instigate a functional change through synergistic conformation events ^13^. However, it does allow us to assess how changing the orientation and inter-unit interactions of one monomer to another along a quasi-similar interface regions alters dipole arrangements. The calculated κ^2^ was 3.79, even closer to the maximal value of 4 than sfGFP^204×2^. While this would suggest an even longer R_0_ distance than GFVen^204^, the inherent function of the GFVen^148^ dimer system makes calculating R_0_ problematic; the donor, sfGFP^SCO148^, is essentially switched off in monomeric state and only becomes activated on dimerisation. However, the main effect that will influence any FRET analysis is the shift in λ_EM_, which is blue shifted by 10 nm in the GFVen^148^ dimer compared to Venus^WT^ (Figure 7d) when excited at a wavelength corresponding to sfGFP. If single wavelength readings are taken with 530 nm assumed to be the Venus emission maximum, fluorescence emission would be underestimated by up to 35% so impacting on perceived FRET efficiency. While residue 204 is more applicable in terms of understanding proximity and FRET due to the non-perturbative nature of the initial mutations, sfGFP^148×2^ and GFVen^148^ still acts as good examples of how proximity is once again having a significant effect on the spectral properties. It also demonstrates that FP dimerisation are not restricted to a defined interaction configuration but that different inter-FP orientations are available.

## Conclusion

Our ability to construct artificial dimers of FPs coupled with structural analysis has allowed us to look at how proximity can influence two of their key functions: inherent electronic excitation/light emission and communication through energy transfer. With regards to the latter, we can use structural information to predict dipole alignments of two CROs, which is critical to FRET through defining κ^2^. In our case, the arbitrary 0.6667 for the κ^2^ value provides a significant underestimate of the predicted values that impacts on R_0_. There are been several studies to date that measure FRET in artificial constructs whereby FPs are coupled via linker sequences or whole protein domains. However, by linking two FPs together they can no longer freely interact with each other due to, for example, steric hinderance (e.g. when using linker sequences) ^15^ or spatially forced apart (e.g. when linked to whole proteins) ^14^, schematically outlined in Figure 8. Our use of bio-orthogonal chemistry allows broader sampling and stabilisation of mutually compatible FP interfaces (Figure 8c); non-compatible FP surfaces do not form covalent bonds so the interface will not persist ^13^. The most powerful use of FRET is monitoring protein-protein interactions whereby the FPs are fused to separate proteins. It can be argued that most FP fusions will not associate in most FRET experiments. However, as FPs will be attached to partner protein that normally associate, and if the two FPs are in close proximity they may well align or even interact in preferential arrangements (Figure 8d). This in turn can affect dipole alignment and even inherent FP function. Naïve docking of FPs along with empirical evidences highlights FPs tendency to oligomerise, which will be enhanced by local high concentrations. It is thus clear from our work that by placing FPs in close proximity can result in changes in the expected fluorescence behaviour.

**Figure 8.**
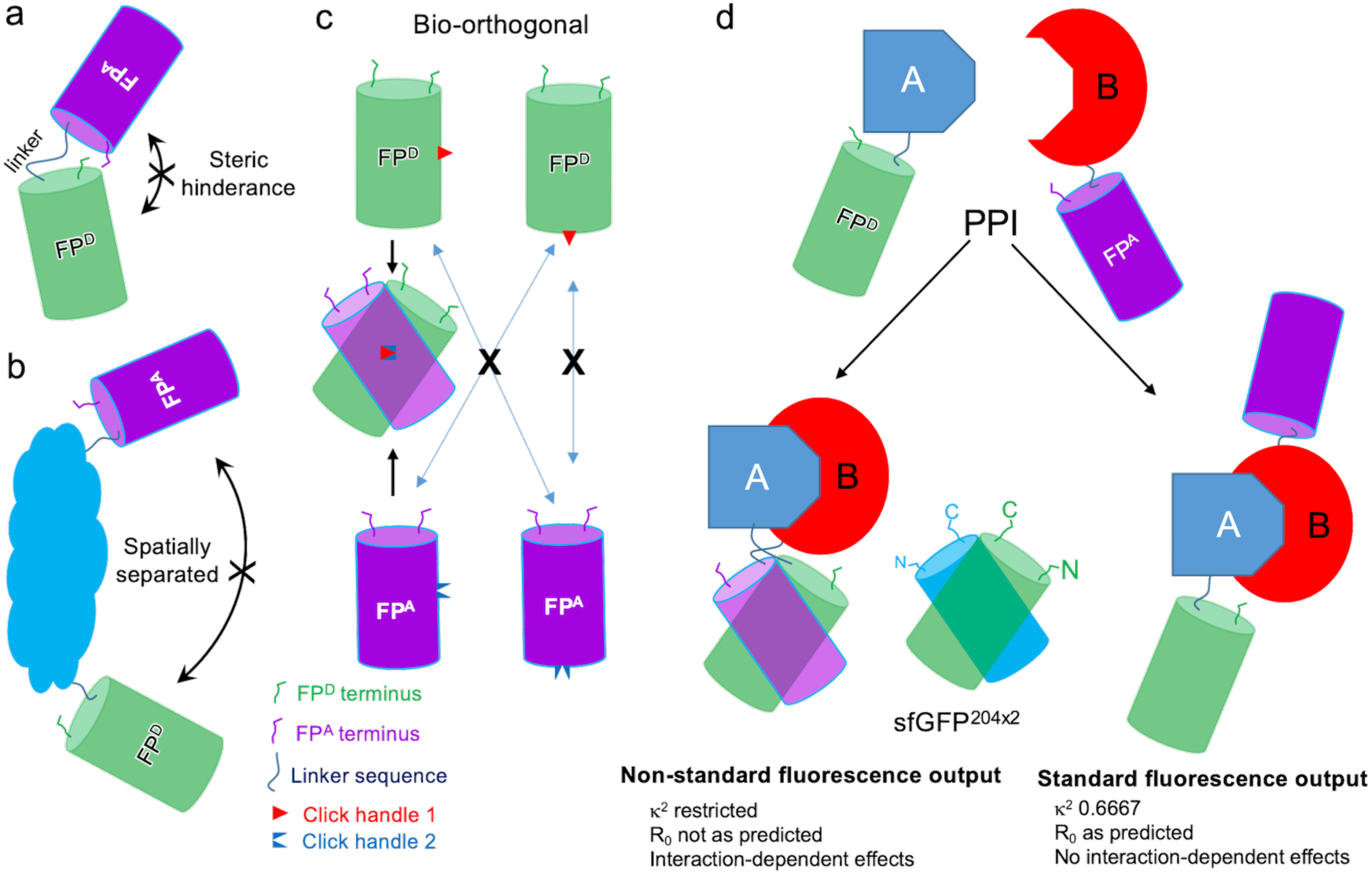
Schematic outline of assessing FP pairs. FP^D^ and FP^A^ refer to nominal FP donor and acceptor for FRET. (a) Model system where short linkers generating a tandem FP pair. Linkers may be too short to allow full freedom to sample interactions, especially side-on interactions. So, there is a low likelihood of FP interaction. (b) Model system whereby FP pairs are bridged by a protein domain or whole protein. If the termini of the bridging protein are at opposite ends then the two FPs will be spatially separated so cannot interact. (c) Bio-orthogonal Click chemistry approach whereby one FP has one type of chemistry (e.g. azide) and the second a mutually reactive handle (shown as red triangle and blue inverted triangle). Only FPs with mutually compatible interfaces will react and so stabilise the interaction. If the interface is not compatible the FPs will not click together. Broader FP-FP interface regions can be sampled through this approach. (d) Protein-protein interaction (PPI) system. Two scenarios are envisaged. The first is that on interaction of protein A and B, the FPs are brought in close proximity to each other promoting association, which may in turn lead to non-standard fluorescence properties. In the second scenario, protein A and B interact but the FPs remain spatially separate so displaying more classical fluorescence behaviour.

## Methods

### Protein production

The monomeric sfGFP^204azF^ and sfGFP^204SCO^ proteins and the sfGFP^204×2^ dimer were produced as described previously ^13^. The WT mCherry and mCherry^198azF^ proteins were produced as outlined in the Supporting Methods.

### Protein dimerisation and conjugation

The procedures for generating sfGFP homodimers and sfGFP-Venus heterodimers have been reported previously ^13^. Generation of the sfGFP-mCherry dimers was performed as follows. The sfGFP^204SCO^ and mCherry^198azF^ were mixed at a equimolar concentration (50 μM, 50 mM Tris-HCl pH 8.0) and left at room temperature for ∼16 hr. Dimers were purified by size exclusion chromatography (Superdex 200, 16/600) and protein concentration determined, as described above. Protein dimerisation and separation was also monitored by SDS PAGE gel. Conjugation with non-proteinaceous molecules is described in the Supporting Methods.

### Steady state absorbance and fluorescence analysis

Spectrophotometry and fluorescence were performed essentially as described previously for sfGFP monomers and dimers, Venus monomers and sfGFP-Venus hybrid dimers ^13^. Analysis of variants involving mCherry followed a similar analysis procedure, using proteins concentration of 5 μM in 50 mM Tris-HCl pH 8.0. Absorbance spectra were recorded on a Cary Win UV, using a 300 nm/min scan rate at 1 nm intervals. Absorbance at λ_max_ for each variant, was used to determine the molar extinction coefficients (ε) for each variant, using the Beer-Lambert equation and measured protein concentrations. Emission spectra were collected on a Cary Varian fluorimeter at a scan rate of 60 nm/min and 1 nm intervals. Emission and excitation slit widths were set to 10 nm and a detector voltage of Low. Samples were excited at 5 nm from 460nm to 590nm as stated in the main text and emission was scanned from the excitation wavelength to 800 nm. J coupling constants (*J*(λ)) were calculated using either available parameters on FPbase ^27^ via the FRET tool or calculated from experimental data using a|e software (http://www.fluortools.com/software/ae). FRET efficiency was calculated using Equation 3.

### Single molecule fluorescence

Measurement and analysis of single molecule sfGFP^204×2^ fluorescence by total internal resonance fluorescence microscopy was performed as described previously ^13^.

### Structure determination of sfGFP^204×2^

The sfGFP^204×2^ dimer variant was concentrated in 50 mM Tris pH 8.0 to a final concentration of 10 mg/ml, and used to set up vapour diffusion crystal trays. A JBScreen membrane (Jena Bioscience, Germany) was used initially to facilitate crystal growth, where large green crystals grew in a multitude of buffer conditions. Large green crystals grew in 20% polyethylene glycol w/v, 100 mM HEPES, which were harvested and transferred to mother liquor supplemented with 13% (w/v) PEG 200 as a cryo-protectant, and vitrified in liquid nitrogen. X-ray scattering data was collected at the Diamond light source, Harwell, UK (beamline IO2). Structure refinement was performed using the CCP4 program suite ^35^. The structure was solved initially using the molecular replacement program PHASER^36^, with wt sfGFP (PDB accession 2B3P) used as a model. Structures were manually adjusted using with COOT^37^, and refined with TLS restrained refinement using REFMAC^38^.

### Kappa^2^ calculation

The dipole orientation factor, κ^2^, was calculated using an approach as described previously ^39^. The model structure of GFVen^204^ dimer was built by overlapping the WT structure of Venus (1MYW) onto the sfGFP^204azF^ component of sfGFP^204×2^ structure. While Venus and sfGFP used here have 15 amino acid differences in the core β-barrel structure only Ala206 in sfGFP^SCO204^ and Val206 in Venus contribute to the domain interface. Residue 204 was then replaced with azF and linked to SCO using the PyMOL bond building tool. The GFVen^204^ model overlaid with sfGFP^204×2^ is shown in Supporting Figure S4. Using the model structure the transition dipole moment (TDM) was calculated using the distance between the centres of the donor and acceptor dyes, and the orientations of the transition dipole moments of the donor, and the acceptor ^39^. The atomic positions of CG2 and C2 of the chromophore, as shown in Supporting Figure S9, were used to the define the vector for the TDM for both Venus and sfGFP ^25^. The κ^2^ was then used in equation 1 together with available experimental to calculate R_0_ for FRET pairs.

## Acknowledgements

We would like to thank the staff at the Diamond Light Source (Harwell, UK) for the supply of facilities and beam time, especially Beamline I03 and I04 staff. We would also like to thank Edward Lemke and his group at EMBL Heidelberg for donating the pEVOL-SCO plasmid. We thank BBSRC (BB/H003746/1 and BB/M000249/1), EPSRC (EP/J015318/1) and Cardiff SynBio Initiative/SynBioCite for supporting this work. RLJ was supported by a Knowledge Economy Skills Scholarship (KESS2) PhD studentship, WDJ by Wellcome ISSF, HLW. by a BBSRC-facing Cardiff University PhD studentship, HSA. by the Higher Committee for Education Development in Iraq. We would like to thank the Protein Technology Hub, School of Biosciences, Cardiff University for use of facilities. We also thank Dr Joachim Goedhart from the University of Amsterdam for his helpful and insightful comments on the manuscript and providing the scripts for calculating fluorescence lifetimes form molar absorbance coefficients.

