## Supporting Methods, Figures and Tables for "The effect of proximity on the function and energy transfer capability of fluorescent protein pairs"

.

Jacob R. Pope[1], Rachel L. Johnson[1], W. David Jamieson[3], Harley L Worthy[1,a], Senthilkumar Kailasam[5,6], Husam Sabah Auhim[1], Daniel W. Watkins[1,b], Pierre Rizkallah[2], Oliver Castell[3], D. Dafydd Jones[1]*.

**Supporting Methods**
**Supporting Tables S1-S2**
**Supporting Figures S1-S10**

**Construction and production of mCherry variants.** The gene encoding wt mCherry resident within the pBAD plasmid was used to generate the K198TAG mutation by whole plasmid site-directed mutagenesis (Forward primer 5'-CGGCGCCTACAACGTCAACATC**TAG**T-3' and reserve primer 5'-GGCAGCTGCACGGGCTTCTT-3') using Q5 polymerase (NEB, USA). *E. coli* Top10 cells were transformed and used to inoculate 1L flasks of LB media supplemented with 100 µg/mL ampicillin, 25 µg/mL tetracycline. To incorporate azF, cells were also co-transformed with pDULEcyanoRS and cultured in the presence of 0.1 mM azF. Cultures were grown for 1 hour at 37°C in a shaking incubator before expression was induced by addition of 0.1% of arabinose and incubated for 24 hours at 25°C. Cultures were kept in the dark until after dimerisation with SCO-K containing variants.

Cells were harvested via centrifugation at 5000 x *g* for 20 mins. The supernatant was discarded, and cells resuspended in 20 mL of 50 mM Tris-HCl pH8.0, 1 mM EDTA. The cells were lysed using a French press and the resulting lysate was clarified by centrifugation at 20,000 xg for at least 30 minutes. Cell lysates were then loaded onto a 5 mL HisTrapHP™ (GE Healthcare) equilibrated in lysis buffer. Bound GFP was eluted by washing the column in 250 mM Imidazole. Samples were then loaded onto a Superdex 75 column equilibrated in 50 mM Tris-HCl pH8.0 and purity was checked via SDS-PAGE analysis. Concentrations of monomer variants were determined using the Bio-RAD DC Protein Assay using wild type wt mCherry as a standard and correlated to the 280 nm absorbance. The quantum of mCherry$^{198azF}$ was calculated as described previously [1] using WT mCherry as the reference sample.

**Conjugation of mCherry with non-proteinaceous molecules.** Conjugation of mCherry$^{198azF}$ with the ncAA SCO-K was performed in 50 mM Tris-HCl pH 8.0) with 5 µM protein and 200 µM of ncAA. Samples were left for 4 hr at room temperature and analysed by absorbance and fluorescence spectroscopy. Conjugation of mCherry$^{198azF}$ with Cy3 DBCO (Click Chemistry Tools, USA) was performed with equimolar concentrations of protein and dye (5 µM). The absorbance and fluorescence emission were recorded immediately after sample mixing. The sample was then left at room temperature overnight. Following overnight incubation, the absorbance and fluorescence were performed again, and the reaction mix was run on SDS page gel.

**Supporting Table S1.** Crystallographic statistics for $sfGFP^{204x2}$

| | **$GFP^{204x2}$** |
|---|---|
| PDB ID | 5NI3 |
| Wavelength (Å) | 0.979 |
| Beamline | Diamond IO2 |
| Space group | $P2_12_12_1$ |
| a (Å) | 96.700 |
| b (Å) | 98.020 |
| c (Å) | 102.640 |
| Resolution range (Å) | 57.17-1.28 |
| Total reflections measured | 1,840,570 |
| Unique reflections | 249,821 |
| Completeness (%) (last shell) | 99.9(99.8) |
| Multiplicity (last shell) | 7.2 (7.1) |
| I/σ (last shell) | 17.7 (1.35) |
| CC1/2 | 0.999 (0.512) |
| R(merge)[a] (%) (last shell) | 4.5(1.68) |
| B(iso) from Wilson ($Å^2$) | 15.12 |
| B(iso) from refinement | 23.7 |
| Log Likelihood Coordinate rms | 0.036 |
| Non-H atoms | 8,482 |
| Solvent molecules | 999 |
| R-factor[b] (%) | 15.8 |
| R-free[c] (%) | 17.3 |
| Rmsd bond lengths (Å) | 0.019 |
| Rmsd bond angles (°) | 2.143 |
| Core region (%) | 98.40 |
| Allowed region (%) | 1.13 |
| Additionally allowed region (%) | 0 |
| Disallowed Region (%) | 0.45 |
| **Structural comparison** | |
| RMSD of azF monomer to sfGFP (Å) | 0.193 |
| RMSD of SCO monomer to sfGFP (Å) | 0.165 |

**Table S2.** Calculated $R_0$ values based on available data

| **Data source** | **Donor** | **Acceptor** | ***J*** **($M^{-1}cm^{-1}nm^{4}$)** | **QY (Donor)** | **$R_0$ (Å) $\kappa^2$=0.667** | **$R_0$ (Å) $\kappa^2$=3.59** |
|---|---|---|---|---|---|---|
| FPbase[a] | sfGFP | Venus | $3.03x10^{14}$ | 0.65 | 55.75 | 73.81 |
| Expt[b] | sfGFP | Venus | $3.77x10^{14}$ | 0.65 | 57.87 | 76.62 |
| Expt[b] | $sfGFP^{204SOC}$ | $Venus^{204azF}$ | $3.46x10^{14}$ | 0.66 | 57.21 | 75.74 |

[a] available from the online fluorescent protein resource, FPbase (https://www.fpbase.org); [b] parameters experimentally determined by the authors.

### Supporting Figures.

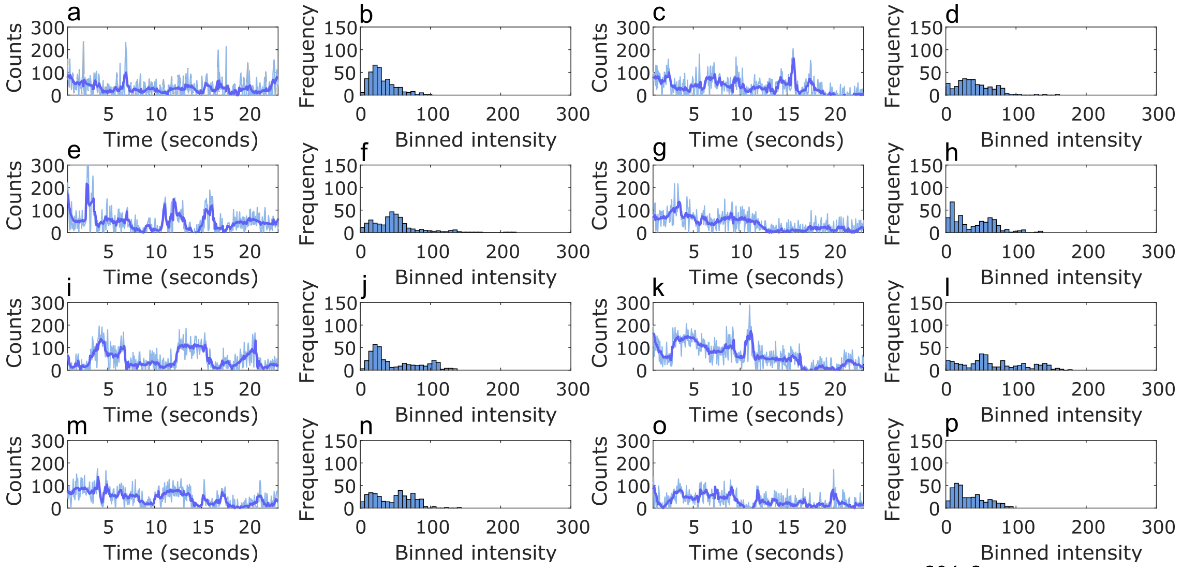

**Supporting Figure S1.** Representative sample of dimeric sfGFP$^{204x2}$ single molecule time course traces (raw and Cheung-Kennedy filtered) coupled with paired intensity frequency histograms (generated from Cheung Kennedy filtered data) to the right of each trace.

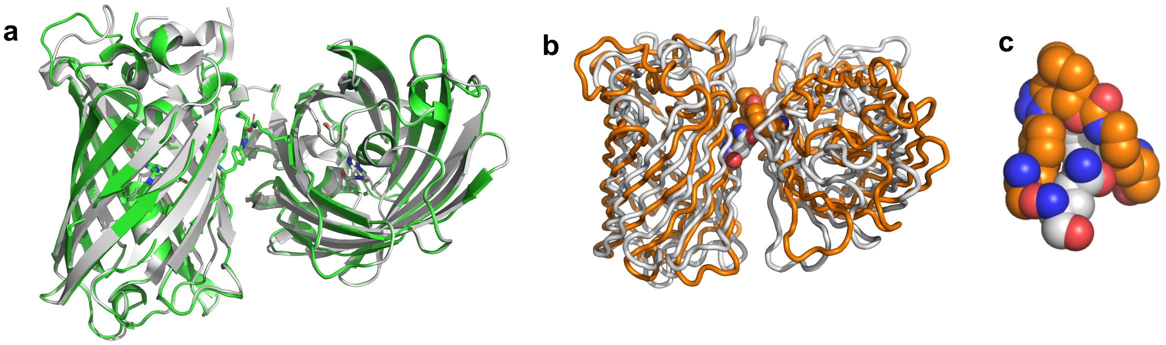

**Supporting Figure S2.** Structural analysis of sfGFP$^{204x2}$. (a) Overlay of the two observed dimers in 5NI3 with the WT sfGFP (grey). The Cα RMSD of WT sfGFP with the azF monomer is 0.193 Å and 0.165 Å with the SCO-K monomer. (b) Overlay of best fitting model calculated previously [2] (coloured grey) and the determined structure of sfGFP$^{204x2}$ (orange). Overlap of residues involved in the triazole crosslink are shown as spheres in (d). The Cα RMSD between the model and determined structure is 5.6 Å.

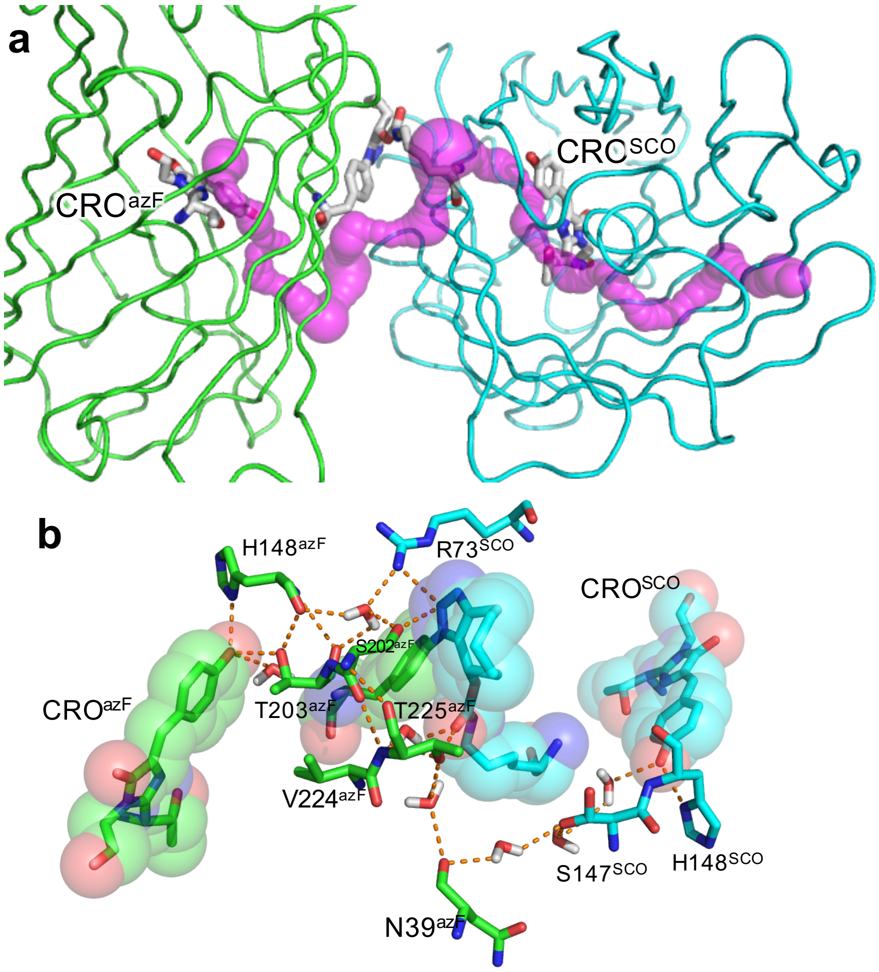

**Supporting Figure S3**. Long range tunnels and bond networks in sfGFP$^{204x2}$. (a) CAVER [3] analysis of sfGFP$^{204x2}$. The tunnel linking the CROs are coloured magenta. (b) Long range H-bond network linking the two CROs. The subscript to the residues denotes the monomer the residues are from. The sfGFP$^{204azF}$ monomer is coloured green and the sfGFP$^{204SCO}$ is coloured cyan. Additional information. Analysis of sfGFP$^{204x2}$ using CAVER 21 suggests that a tunnel linking the two CROs is present (Figure 5a). The pathway extends beyond CRO$^{SCO}$ to exit around residues 23, 54 and 136. An extended putative interaction polar network involving the SCO and triazole moieties, water and amino acids spans the cavity and links the two CROs. Compared to the GFP$^{148x2}$, the network is not as coherent or direct.

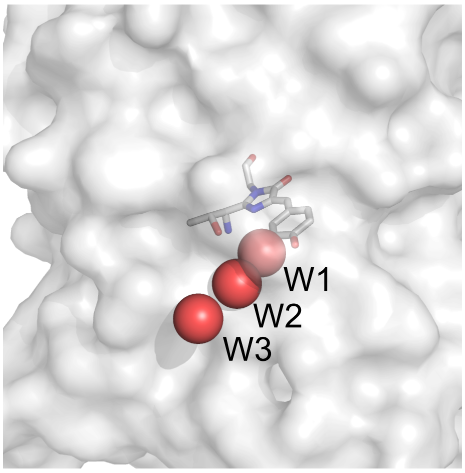

**Supporting Figure S4.** Water molecules (red spheres) closely associated with the CRO (grey sticks) in $sfGFP^{WT}$ monomer.

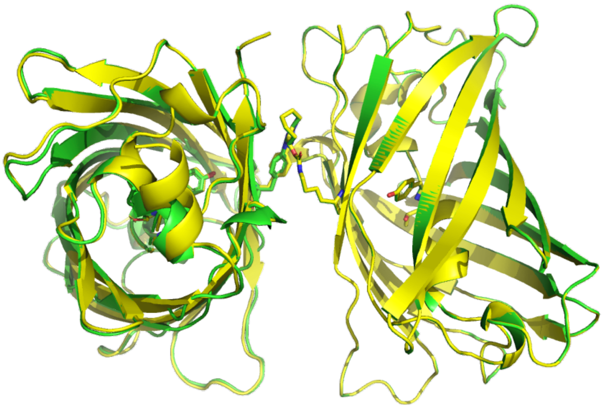

**Supporting Figure S5.** Overlap of $sfGFP^{204x2}$ (green) and model of the $GFVen^{204}$ (yellow). The Cα RMSD between the structure is 0.072 Å.

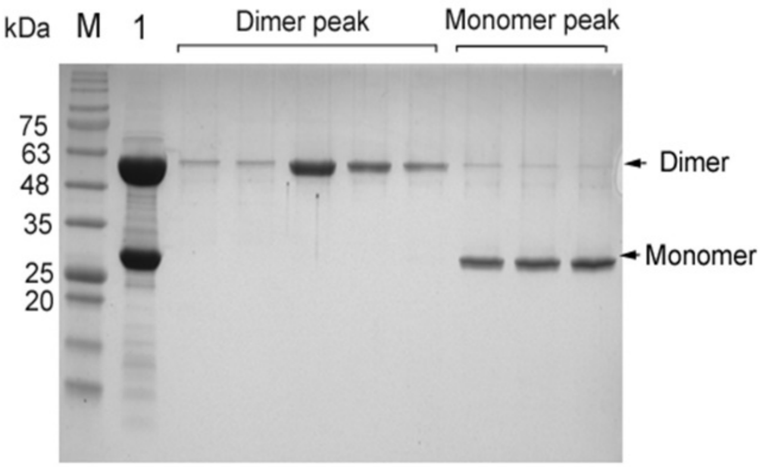

**Supporting Figure S6.** Elution profile of $GFVen^{204}$ dimer and monomeric forms. SDS-PAGE analysis of elution fractions from SEC purification of $GFVen^{204}$ dimer.

Lane M corresponds to the protein marker of known molecular weights. Lane 1 corresponds to the mixture of the click dimerisation reaction before loading onto the SEC column. The dimer corresponds to the first major elution peak and monomer corresponds to the second elution peak on the chromatogram SEC profile.

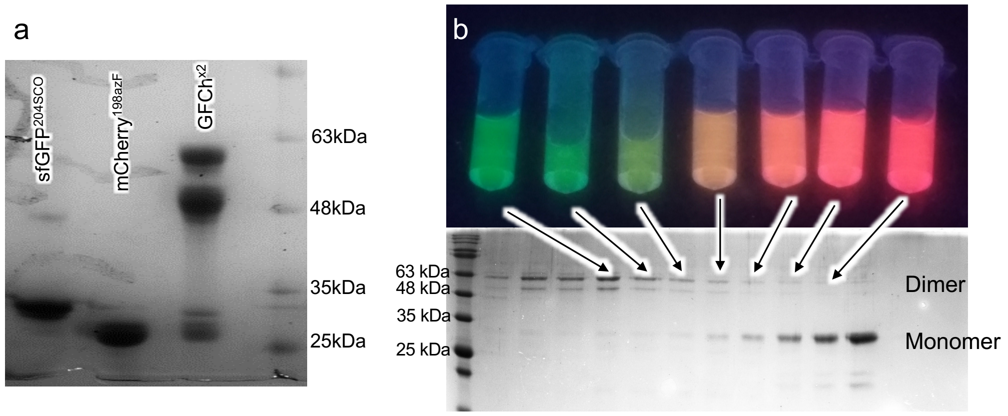

**Supporting Figure S7.** Dimerisation of sfGFP$^{204SCO}$ and mCherry$^{198azF}$. (a) SDS PAGE analysis of the dimerisation reaction. The GFCh$^{x2}$ lane is the endpoint of the reaction. Two bands for mCherry associated proteins have been reported previously by [4], which is due to hydrolysis of the chromophore on boiling. (b) Separation of the dimer from monomer. The top panel represents fluorescence output from the samples equivalent to the fractions analysed by SDS PAGE in the lower panel. On illumination with a transilluminator, the dimer fractions are green, mixed fractions are orange and mCherry based-monomers are magenta is colour. Monomer fractions equivalent to sfGFP$^{204SCO}$ elute after mCherry (not shown).

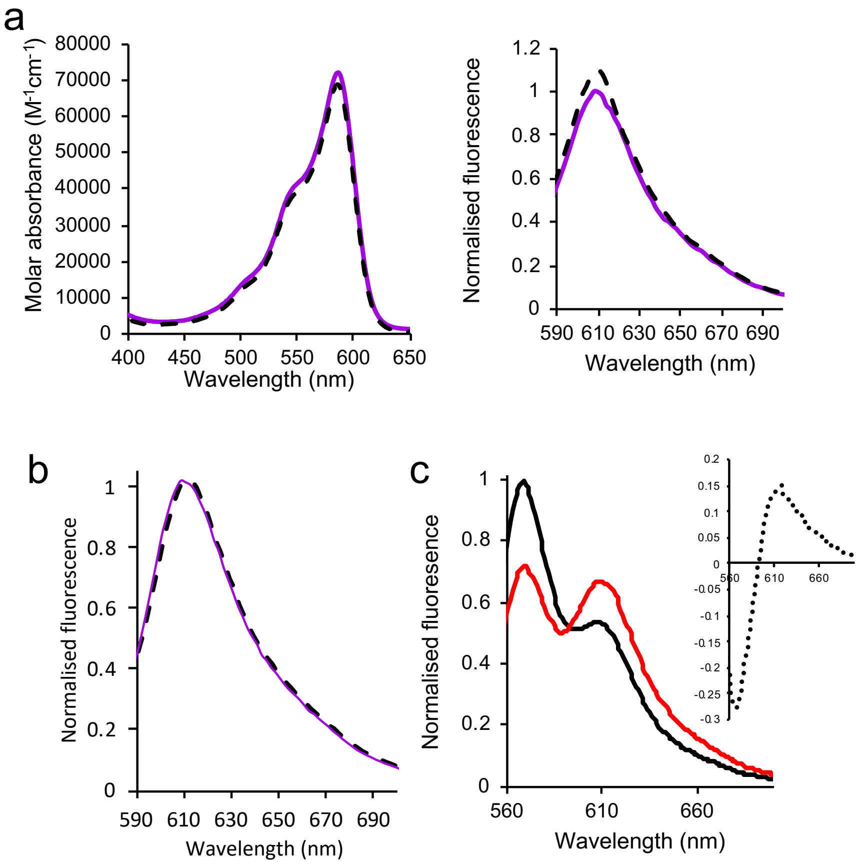

**Supporting Figure S8.** Spectral properties of mCherry$^{198azF}$ and various conjugates. (a) Molar absorbance (left) and fluorescence emission on excitation at 587 nm (right) of WT mCherry (purple) and mCherry$^{198azF}$ (black dashed lines). Fluorescence emission was normalised to WT mCherry. (b) Fluorescence emission (on excitation at 587 nm) on incubation of mCherry$^{198azF}$ with 20 folded excess of SCO-K at the start (0 hr, black dashed lines) and after 4 hr (purple line). Fluorescence normalised to 0 hr time point. (c) Fluorescence emission (on excitation at 555 nm) on incubation of mCherry$^{198azF}$ with DBCO-Cy3 for 0 hr (black line) and overnight (red line). Fluorescence is normalised to the 0 hr time point. Inset the subtraction spectra of the overnight sample from the 0 hr time point. The estimated *J* coupling according to FPbase between Cy3 (donor) and mCherry (acceptor) is $4.83 \times 10^{15}$ $M^{-1}cm^{-1}nm^4$.

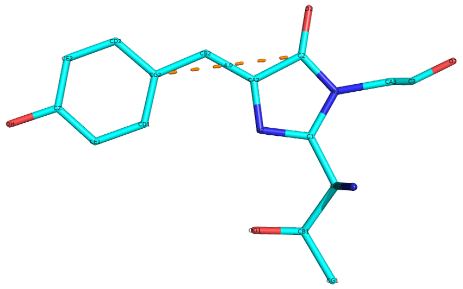

**Supporting Figure S9.** Atomic positions in the chromophore used to define the vector the transition dipole moment for calculating $\kappa^2$

1. Reddington, S.C. et al. Genetically encoded phenyl azide photochemistry drives positive and negative functional modulation of a red fluorescent protein. *RSC Advances* **5**, 77734-77738 (2015).
2. Worthy, H.L. et al. Positive functional synergy of structurally integrated artificial protein dimers assembled by Click chemistry. *Communications Chemistry* **2**, 83 (2019).
3. Chovancova, E. et al. CAVER 3.0: a tool for the analysis of transport pathways in dynamic protein structures. *PLoS Comput Biol* **8**, e1002708 (2012).
4. Gross, L.A., Baird, G.S., Hoffman, R.C., Baldridge, K.K. & Tsien, R.Y. The structure of the chromophore within DsRed, a red fluorescent protein from coral. *Proc Natl Acad Sci U S A* **97**, 11990-5 (2000).
